# Specific mutations in genes responsible for Alzheimer and for Duchenne Muscular Dystrophy introduced by Base editing and PRIME editing

**DOI:** 10.1101/2020.07.31.230565

**Authors:** Joël Rousseau, Cédric Happi Mbakam, Antoine Guyon, Guillaume Tremblay, Francis Gabriel Begin, Jacques P. Tremblay

## Abstract

Base editing technique and PRIME editing techniques derived from the CRISPR/Cas9 discovery permit to modify selected nucleotides. We initially used the base editing technique to introduce in the *APP* gene the A673T mutation, which prevents the development of Alzheimer’s disease. Although the desired cytidine to thymidine mutation was inserted in up to 17% of the *APP* gene in HEK393T, there were also modifications of up to 20% of other nearby cytidines. More specific mutations of the *APP* gene were obtained with the PRIME editing technique. However, the best percentage of mutations was only 5.8%. The efficiency of the PRIME editing treatment was initially tested on the *EMX1* gene. A single treatment produced the desired modification in 36% of the *EMX1* gene. Three consecutive treatments increased the percentage of mutations to 50%. The PRIME editing technique was also used to insert specific point mutations in exons 9 and 35 of the *DMD* gene coding for the dystrophin gene and which is mutated in Duchenne Muscular Dystrophy (DMD). Up to 10% desired mutations of the *DMD* gene were obtained. Three repeated treatments increased the percentage of specific mutations to 16%. Given that there are thousands of nuclei inside a human muscle fiber and that the dystrophin nuclear domain is about 500 μm, this level of modifications would be sufficient to produce a phenotype improvement in DMD patients.

## INTRODUCTION

The rapid development of gene editing techniques derived from the CRISPR/Cas9 discovery now permits to modify the human genome and opens the possibility of developing therapies for hereditary diseases. In particular, the base editing technique^1^ and the PRIME editing technique^2^ permit to modify a single targeted nucleotide. The base editing technology ^1^ uses a Cas9 nickase (D10A), permitting to nick the non-edited strand, fused with a cytidine deaminase to chemically modify a cytidine (C) into a thymine (T) in a window of 5 nucleotides included in the protospacer sequence. This targeted sequence is called the Protospacer sequence. The PRIME editing technology uses a Cas9 nickase (H840A), permitting to nick the edited stand. The pegRNA includes the sgRNA variable and constant sequences prolongated by nucleotide sequences coding for the Reverse Transcriptase Template (RTT) and a Primer Binding Site (PBS). The RTT contains nucleotides, which are complementary to the sense strand except for the nucleotide(s) to be modified. In the present article, we have used these techniques to modify the human Amyloïd Precursor Protein (*APP*) gene and the *DMD* gene. Mutations of the *APP* gene are either responsible for familial forms of Alzheimer disease^3^ or may prevent the development of this disease ^4^. Mutations of the *DMD* gene coding for dystrophin are responsible for Duchenne Muscular Dystrophy^5^.

The APP protein is normally cut by the α-secretase followed by a cut by the γ-secretase. This degradation pathway does not produce any pathological peptide. However, the APP protein may also be first cut by the β-secretase followed by the γ-secretase resulting in the production of the β-Amyloid peptide (Aβ). These Aβ peptides aggregate to each other producing amyloid plaques responsible for neuron death and memory problem in Alzheimer patients. Jonsson et al. have shown in 2012 ^4^ that the A673T mutation in the APP protein located near the β-secretase cut site protects again the development of Alzheimer even in person that are more than 95 years old. The first experiments presented in this article demonstrated that it is possible to introduce by base editing and by PRIME editing this specific point mutation in the human *APP* gene.

About 30% of the Duchenne Muscular Dystrophy (*DMD*) cases are due to point mutations in one of the 79 exons present in that gene ^5^. Most of these point mutations are single nucleotide mutations resulting in a nonsense codon. In the present article, we have shown that the PRIME editing technology can produce specific point mutations in the *DMD* gene.

## RESULTS

### Mutation of the Amyloïd Precursor Protein (*APP*) gene

#### Mutation of the *APP* gene induced by base editing

Our first experiment aimed to introduce the A673T mutation in the *APP* gene. We thus aimed to change the alanine codon (5’GCA3’) in position 673 into a threonine codon (5’ACA3’). We initially aimed to introduce this specific mutation in the *APP* gene using the base editing technology^1^.We targeted the antisense sequence (5’TGC3’) of the alanine codon to modify the cytidine into a thymine thus producing the antisense sequence (5’TGT3’) of the threonine codon (Figure 1A). The base editing technique requires the presence of a Protospacer Adjacent Motif (PAM) located at about 15 to 20 nucleotides in the 3’ direction from the targeted nucleotide and permits to edit the cytidines located within a 5 nucleotides window at the 5’ end of the protospacer sequence targeted by the sgRNA. There was no adequate SpCas9 PAM in an adequate position in the antisense strand. However, an adequate NGAN PAM was present for the genetically modified SpCas9_VQR_ nickase (SpCas9n_VQR_)^6^ (Figure 1A). We have thus fused the SpCas9n_-VQR_ nickase gene with the activation induced cytidine deaminase (Target-AID) to produce the SpCas9n_VQR_-AID gene. HEK293T cells were transfected with one plasmid coding for this SpCas9n_VQR_-AID and with a second plasmid coding for the sgRNA under a U6 promoter. The PAM required for the DNA binding of the SpCas9n_VQR_protein is underlined by a tick red line in Figure 1A. Different promoters (EF1 and CBH) were used to control the expression of the SpCas9n_VQR_-AID gene. Three days later the DNA was extracted from the cells and *APP* exon 16 was PCR amplified (primers in Table 1). The amplicons were sent for Sanger sequencing and the sequencing results were analyzed with EditR web site (https://moriaritylab.shinyapps.io/editr_v10/) (Figure 1B). This analysis indicated that depending on the plasmid used to express the SpCas9n_VQR_-AID gene there were 12 to 17% of the targeted C (C2 in Figure 1A) that was modified into a T, thus inducing the desired A673T mutation (Figure 1B). However, the EditR analysis also indicated that other cytidines (C1, C3, C4 and C5, Figure 1A and B) located in or near the editing window were also mutated into a T resulting in the modifications of other amino acids in the APP protein. To verify these undesired results, the exon 16 amplicons were also sent for Illumina deep sequencing. This more exact analysis of 5000 to 10000 amplicons confirmed that 14 to 21% of the targeted C were indeed modified into a T (Figure 1C). However, this second analysis also confirmed that other cytidines were also mutated into a thymine. In particular nucleotide C1 was modified in 17 to 21% of the amplicons. As observed in figure 1C, conversion efficiency depends on the position of the C in the guide. For C3 to C5 located in positions 7 to 11 of the sgRNA, the C to T modification drop drastically. Since the modifications of these other cytidines did result in modifications of other amino acids, we decided to test the newly discovered PRIME editing technology to verify whether it would produce more specific nucleotide mutations.

**Table 1:**
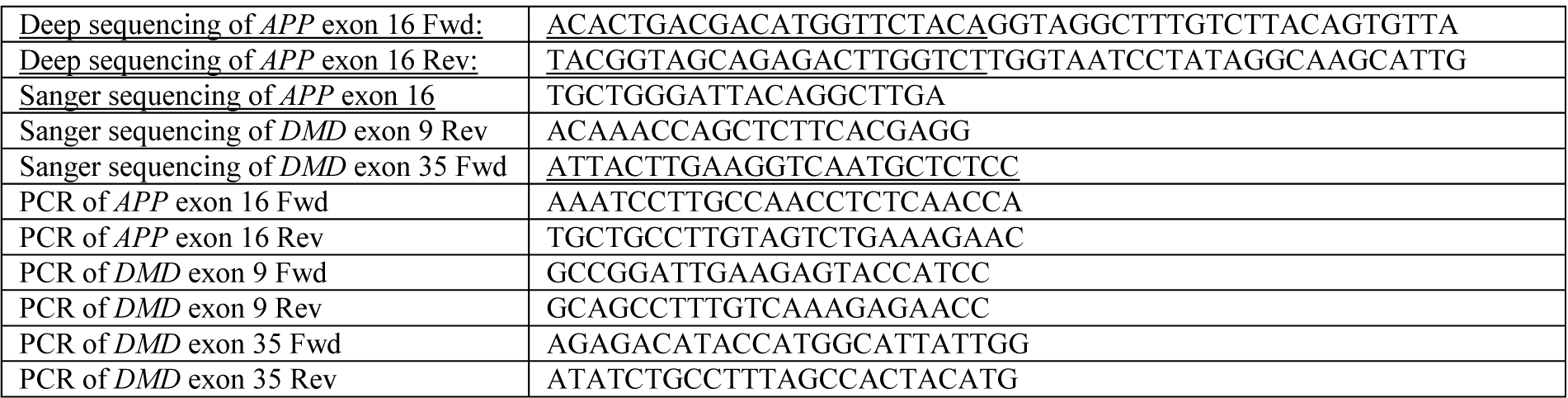
Primer sequence table.

**Figure 1:**
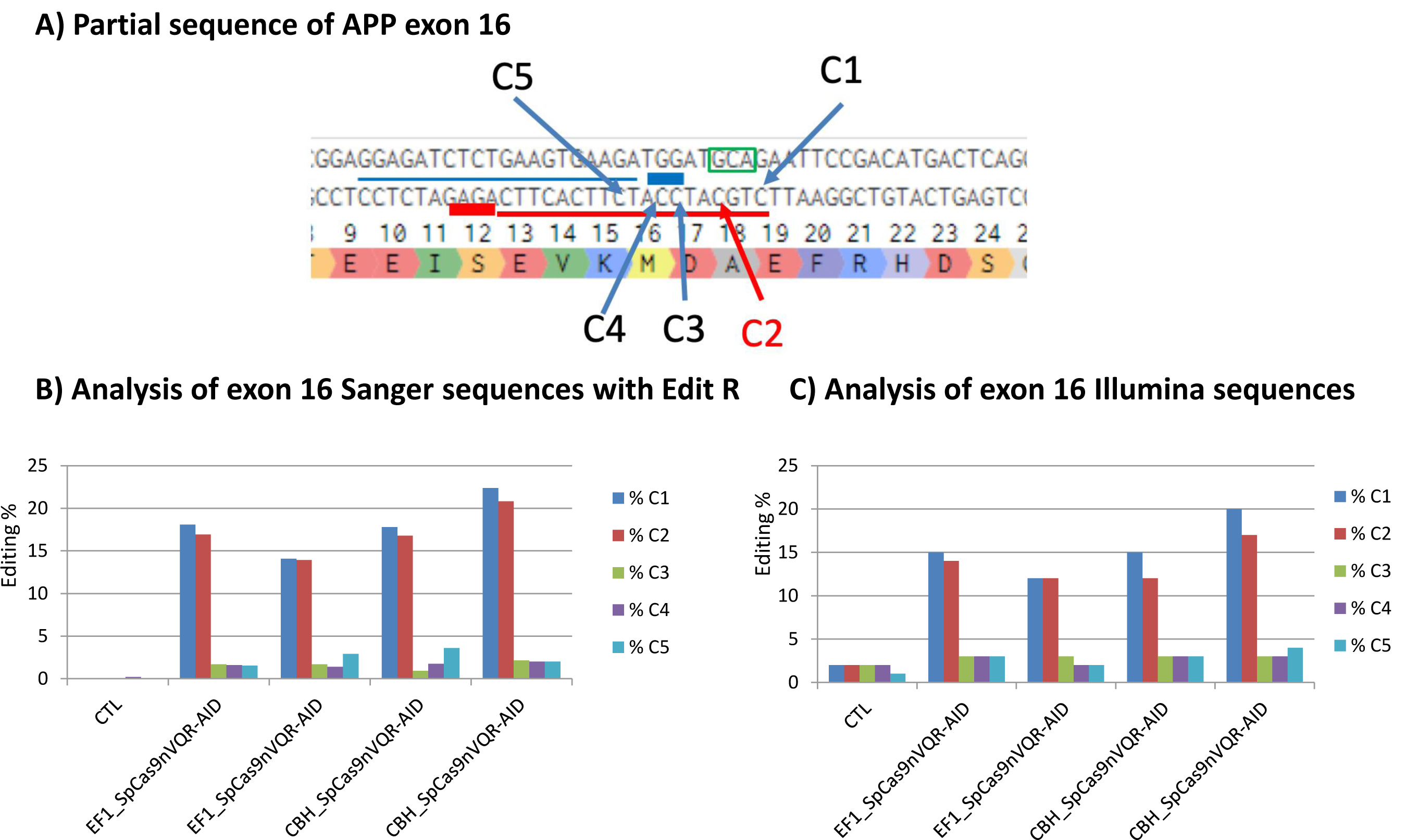
Mutation of the *APP* gene by base editing. **A)** Partial sequence of exon 16 of the *APP* gene. The sequence surrounded by a green box is the alanine codon to be modified into a threonine codon. The sequence (5’AGAG3’) underlined by a thick red line in the antisense strand is the PAM for the SpCas9n_VQR_-AID and the sequence underlined by a thin red line is the protospacer sequence targeted by the sgRNA. There are 5 cytidines (C1 to C5) in the sequence targeted by the sgRNA, which are in the editing window of the SpCas9n_VQR_-AID. The cytidine C2 has to be mutated into a T to induce the desired Alanine to Threonine mutation. The sequence underlined by a thick blue line in the sense strand is the NGG PAM for the SpCas9n_H840A_-reverse transcriptase for PRIME editing and the sequence underlined by a thin blue line is the protospacer sequence targeted by the sgRNA component of the pegRNA. **B)** The Sanger sequencing results were analyzed with the EditR web site (https://moriaritylab.shinyapps.io/editr_v10/) following base editing. The experiment was done in duplicates for each plasmid coding for the SpCas9n_VQR_-AID under the EF1 or the CBH promoter. The EditR analysis indicates that the nucleotide C2 is the most modified by the base editing technique. However, EditR also suggests that cytidines (C1, C3, C4 and C5) present in the antisense strand of the *APP* exon 16 are mutated by the base editing treatment. **C)** Analysis of the amplicon sequences using Illumina deep sequencing confirmed that the nucleotides C2 are mutated. However, the nucleotides C1, C3, C4 and C5 are mutated in less than 0.2% of the amplicons, i.e., at the background level.

#### Mutation of the *APP* gene induced by PRIME editing

We selected a SpCas9nH840Anickase had a PAM near the *APP* 673 codon. We then designed 9 different pegRNAs having different length of Primer Binding Site (PBS) and of Reverse Transcriptase Template (RTT) (see list of these pegRNAs in Figure 2A). We transfected HEK293T cells with the PRIME editor 2 (pCMV-PE2) plasmid obtained from Addgene inc. (Plasmid # 132775). It encodes a Moloney murine leukemia virus reverse transcriptase (M-MLV RT) fused through a flexible linker with the SpCas9 H840A nickase. The cells were simultaneously transfected with a second plasmid encoding a pegRNA under a U6 promoter (pU6-pegRNA-GG-acceptor, Addgene plasmid #132777). DNA was extracted from the cells 3 days later. *APP* exon 16 was then amplified by PCR (Primer sequence in Table 1). The amplicon sequences were obtained by Sanger sequencing because this technique provided a more rapid turnover of the results. The sequencing results were analyzed with the EditR. This web site indicated that depending of the peg configuration, between 2 to 5% of the targeted cytidines C2 (using the same numbering of the C as for the Base editing technique) were modified into a T as desired (Figure 2B). However, the EditR analysis suggested that cytidines C1, C3, C4 and C5 were also modified into a T in about 2% of the amplicons (Figure 2B). This level of potential modifications is the same as the level observed in the untreated control. To verify the editing results more precisely, the exon 16 amplicons were also sequenced using Illumina deep sequencing technology. This more precise analysis of 5000 to 10000 amplicon sequences confirmed that depending on the pegRNA configuration, the C2 cytidine was mutated into a thymine in 0.7% up to 5.8% of the amplicons (Figure 2C). The best results were obtained with the pegRNA1. More importantly, the Illumina deep sequencing results indicated that the C1, C3, C4 and C5 were not modified into a T (Figure 2C). Moreover, the other non-cytidine nucleotides located near the targeted C2 nucleotide were not changed above the background level. Thus, the PRIME editing technology provided a clean modification of the targeted nucleotide.

**Figure 2:**
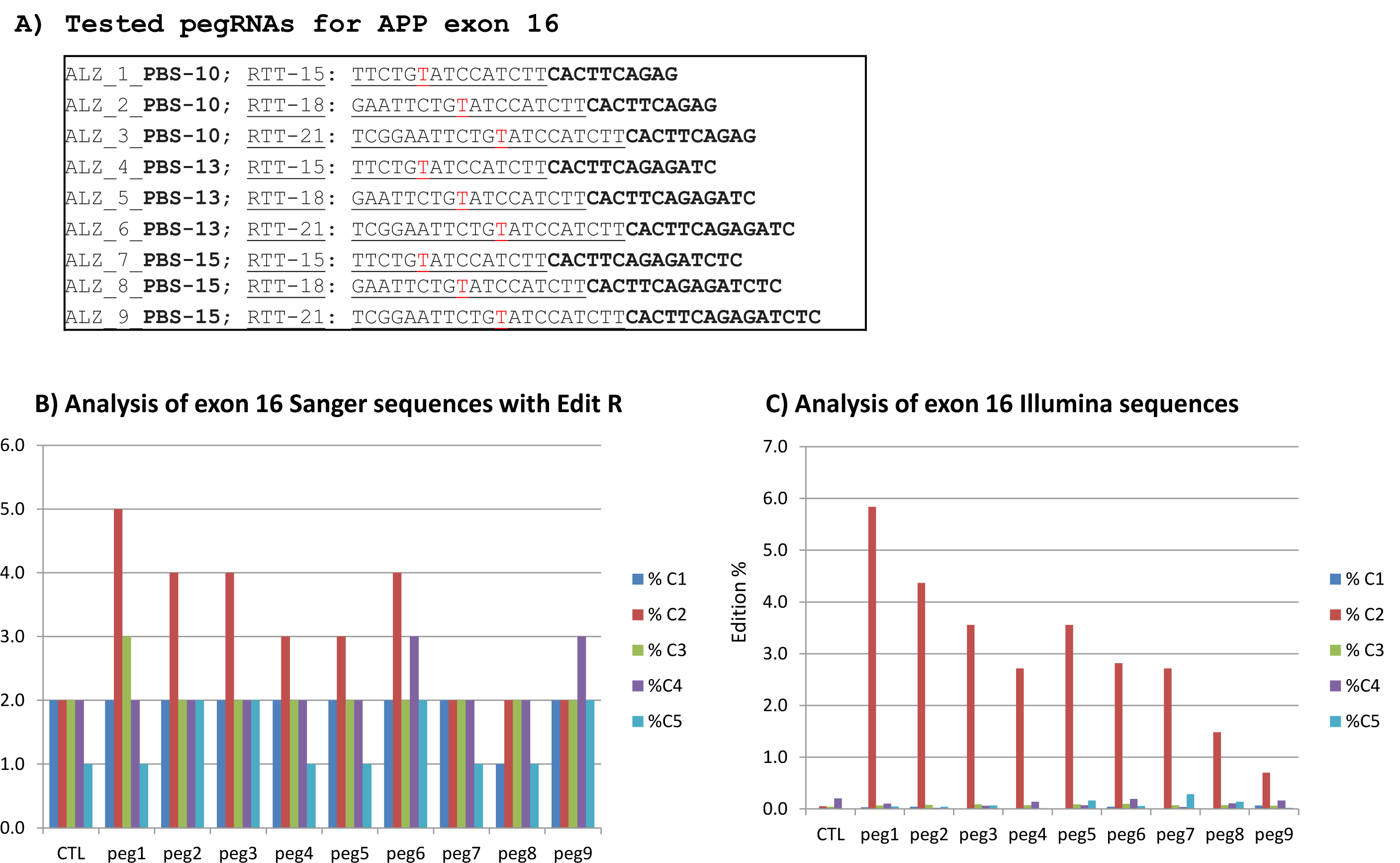
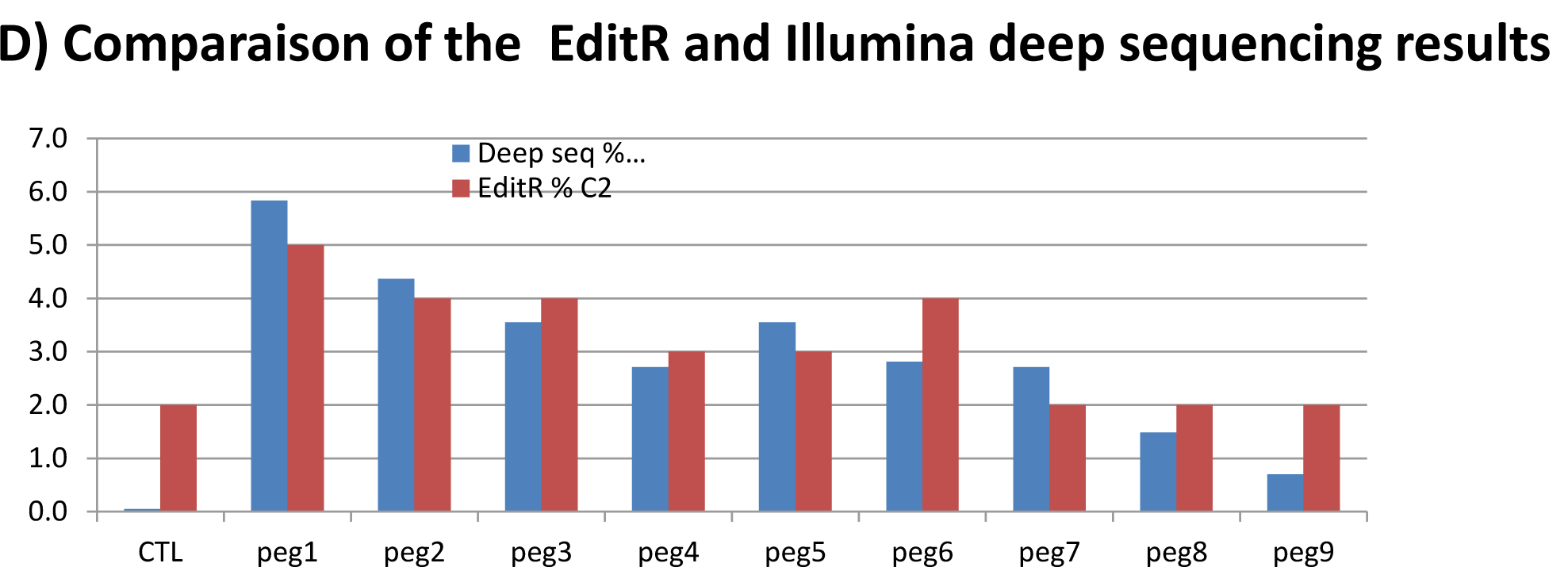
Mutation of the *APP* gene by PRIME editing. **A)** Nine pegRNAs targeting exon 16 of the sense strand of the *APP* were tested. The sequences of the Primer Binding Site (PBS, in bold) and of the Reverse Transcriptase Template (RTT, underlined) of the pegRNAs are listed. The T in red in the RTT permits to introduce an adenosine (A) in the sense strand to mute the alanine (GCA) codon into a threonine (ACA). **B)** HEK293T cells were transfected with plasmid pCMV-PE2 and with a plasmid coding for one of the 9 pegRNAs under a U6 promoter. Three days later, part of exon 16 was amplified by PCR and the amplicon sequences were obtained by Sanger sequencing and analyzed with EditR. The targeted cytidine (C2) is mutated 2 to 5% of the sequences. However, the EditR analysis suggests that 2% of the cytidines C1, C3. C4 and C5 are also mutated. **C)** The amplicon sequences of *APP* exon 16 were also analyzed by Illumina deep sequencing. This analysis confirmed that the C2 nucleotide was mutated in 0.7 to 5.8% of the sequences. However, this analysis indicated that the nucleotides C1, C3, C4 and C5 were not mutated by PRIME editing. **D)** Side by side comparison of the percentages of cytidine C2 mutation determined by the Sanger/EditR analysis and the Illumina deep sequencing analysis. Although imperfect, the EditR analysis permits a preliminary analysis of the results.

The side by side comparison of the EditR analysis and Illumina sequencing results (Figure 2D) indicated that although the analysis of the Sanger sequences with EditR does not provide perfect results, the percentage of mutations detected by this technique is closely parallel to the results obtained with deep sequencing. Thus, the EditR analysis can be used for a rapid screening to detect point mutations produced with the PRIME editing technology.

### Mutation of the Duchenne Muscular Dystrophy (*DMD*) gene by PRIME editing

To confirm that the PRIME editing technology may provide a clean modification of a specific nucleotide, we also tested the modification of a single nucleotide mutation in exon 9 of the *DMD* gene. We selected the SpCas9 H840Anickase because that was a PAM located close to the targeted nucleotide (Figure 3A) and we tested 9 different pegRNAs (sequences in Figure 3B). Since we did not have cells from DMD patients having a nonsense mutation, our initial experiment aimed to mutate a specific nucleotide in a CGA arginine codon to modify it into a TGA stop codon to create cell lines for future experiments. The plasmid pCMV-PE2 (Addgene #132775) coding for the SpCas9 H840A nickase fused with the M-MLV reverse transcriptase (M-MLV RT) and a plasmid coding for one of the pegRNAs under a U6 promoter were transfected in HEK293T cells. The sequence in the RTT of the different tested pegRNAs (sequence list in Figure 3B) permitted to produce the intended mutation. A pegRNA used in Anzalone et al article ^2^ to mutate a nucleotide in the EMX1 gene was also used as a positive control. DNA was extracted from the cells 3 days later. The percentage of modifications of the targeted cytidine into a thymine was analysed only by Sanger sequencing and EditR analysis. The pegRNA targeting the EMX1 gene produced a 36% (Figure 3C) or 33% (Figure 4C) modification of the targeted nucleotide. A negative control transfected only with the pU6-pegRNA-GG-acceptor, the EditR analysis suggested that there was a 6% background presence of a T nucleotide in the position of the targeted C (Figure 3C). This is probably due to the background noise of the Sanger sequencing or the heterogeneity of the cell population for that gene sequence. However, when the cells were transfected with the pCMV-PE2 and one of the 9 pegRNAs encoded in plasmid pU6-pegRNA-GG-acceptor (Addgene #132777), there was 6 to 10% of the targeted cytidine that were a thymine (Figure 3C. Given the background noise, this suggests that there were about 4% real mutations of the targeted cytidine into a thymine by pegRNAs 2, 4, 5 and 9.

**Figure 3:**
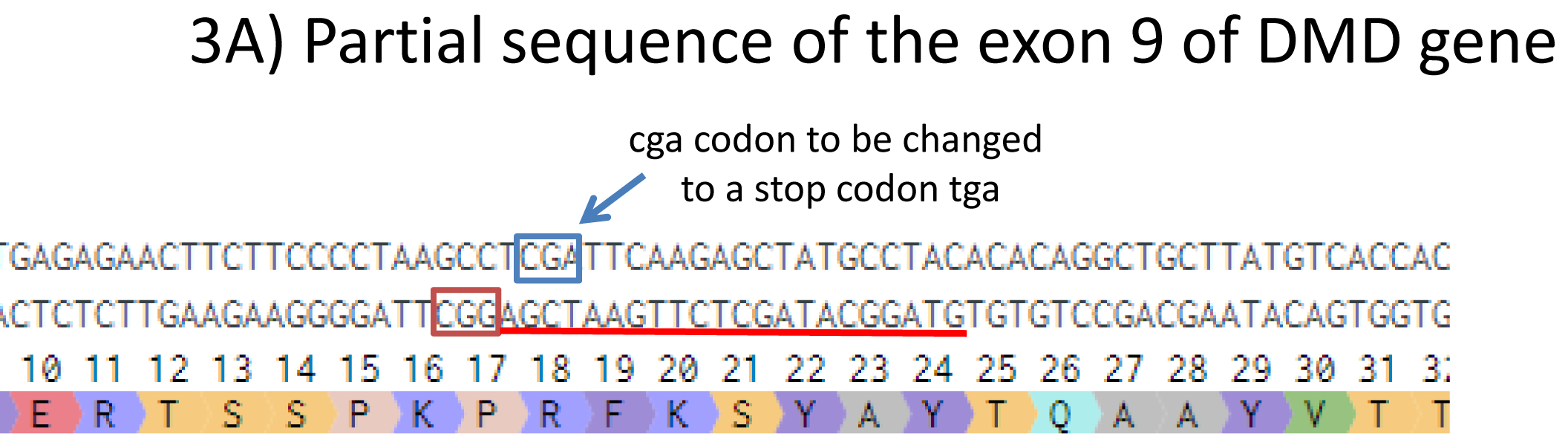
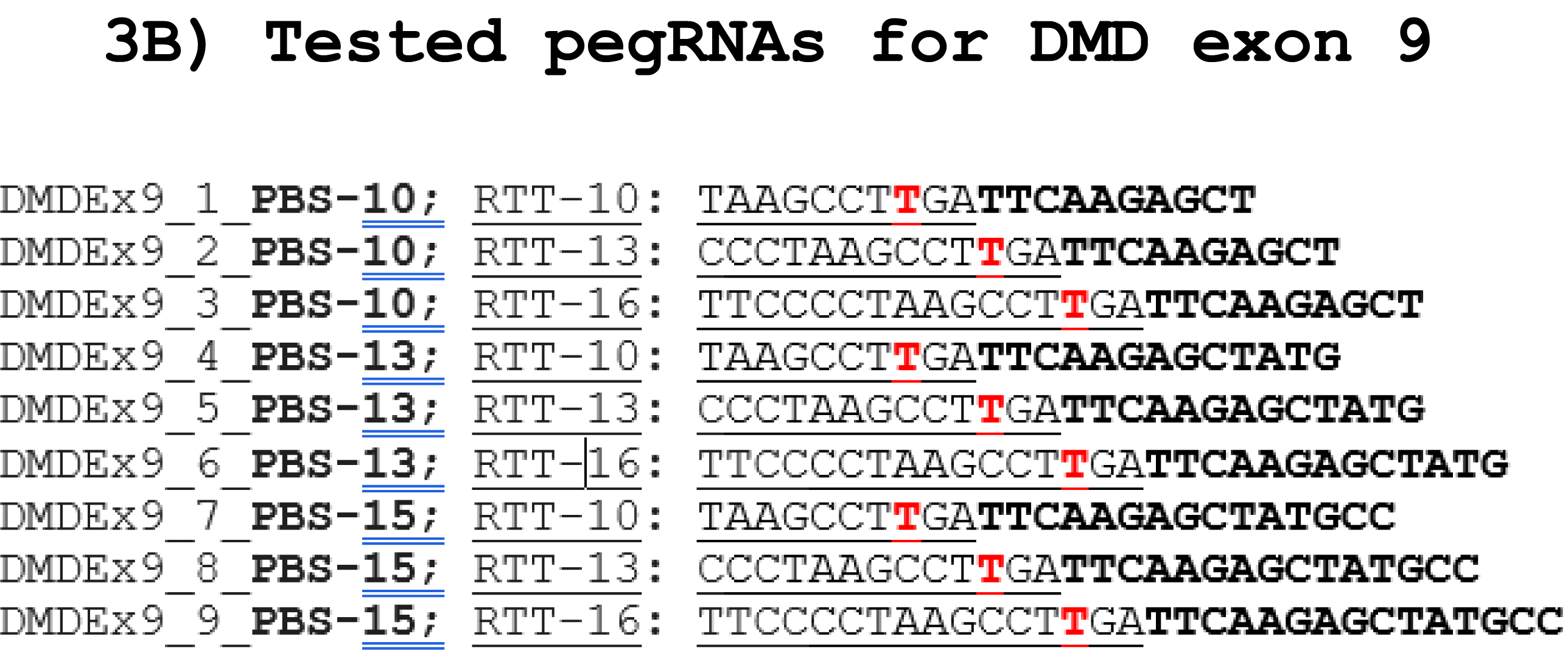
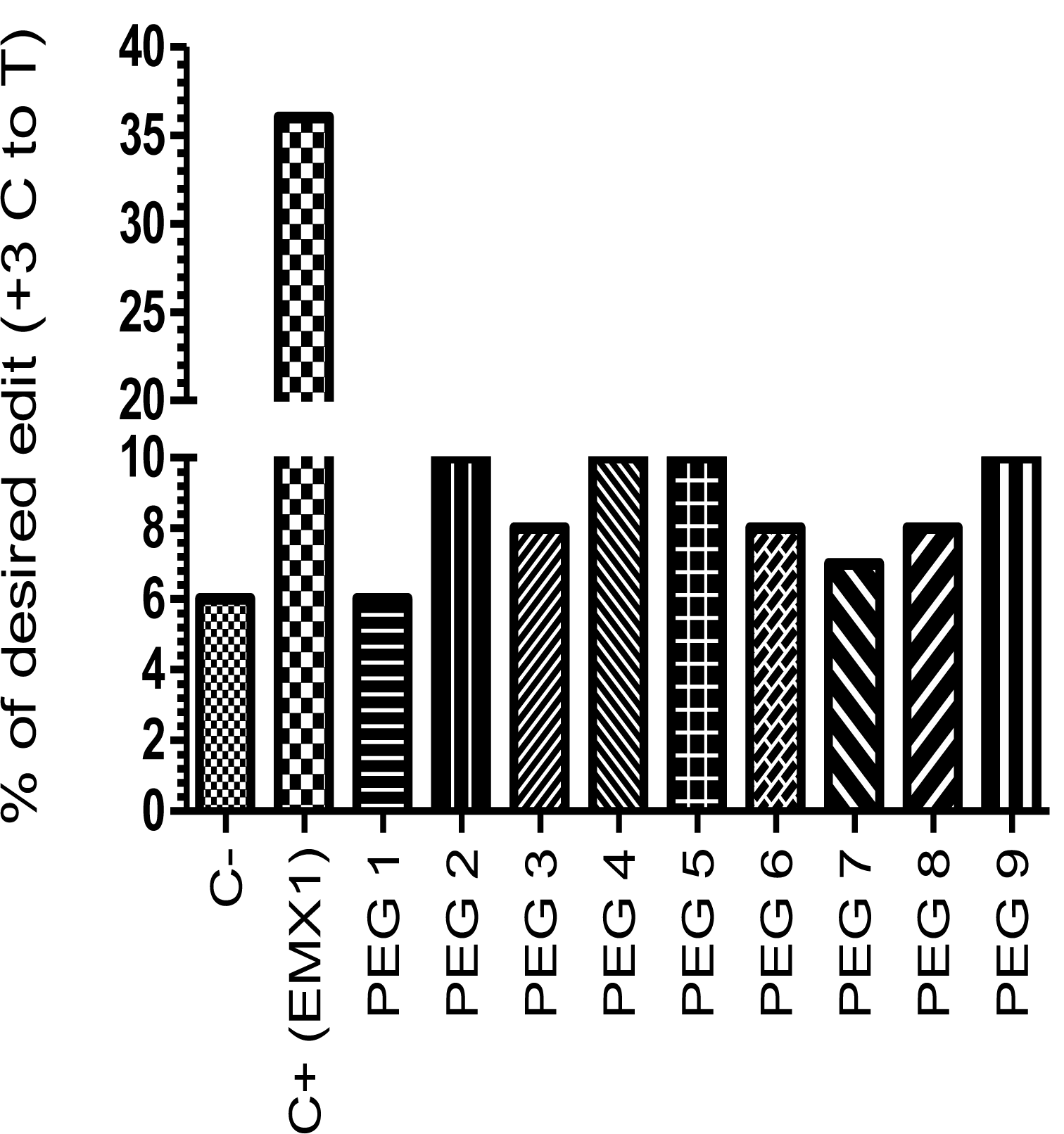
Prime editing of *DMD* exon 9. **A)** Partial sequence of exon 9 of the *DMD* gene. The SpCas9n PAM is in a red box and of the pegRNA protospacer sequence is underlined in red. **B)** The sequences of the Primer Binding Site (PBS, in bold) and of the Reverse Transcriptase Template (RTT, underlined) of 9 pegRNAs targeting *DMD* exon 9. The T in red in the RTT permits to introduce an adenosine (A) in the strand containing the PAM (in the present case the antisense strand). This results in the presence of a thymine in the sense strand modifying the arginine codon into a stop codon. **C)** Results of Sanger sequencing and EditR analysis of PRIME editing of the *DMD* exon 9. There is a high background level (6%) of thymidine (T) in the position of the targeted nucleotide. However, pegRNAs 2 to 9 increased the percentage of T up to 10%. A control pegRNA targeting the EMX1 gene produced a 36% modification in that gene.

**Figure 4).**
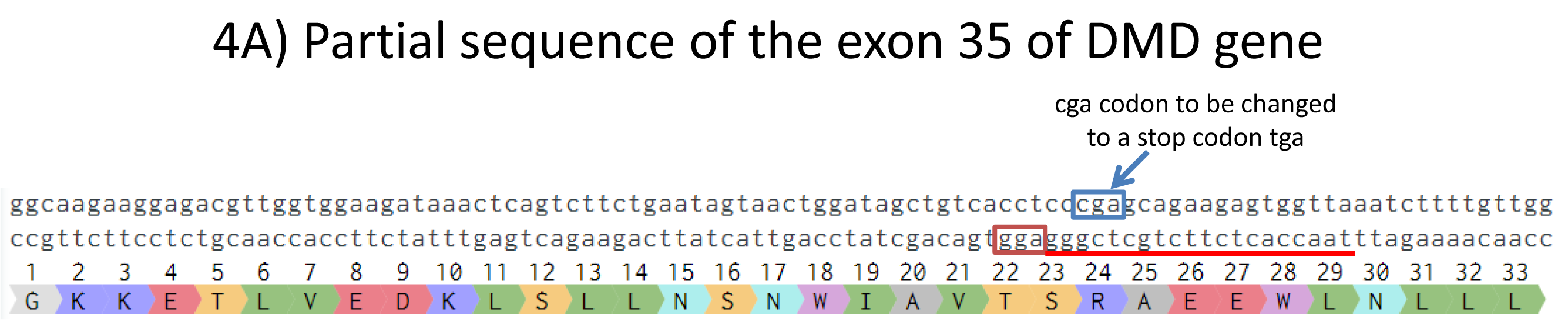
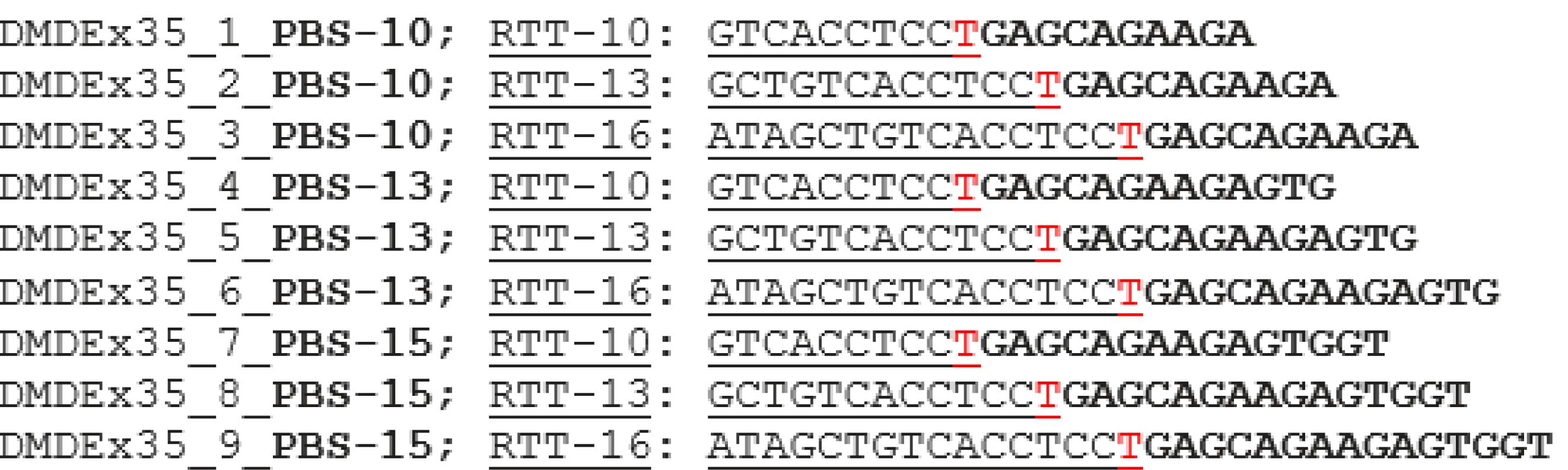
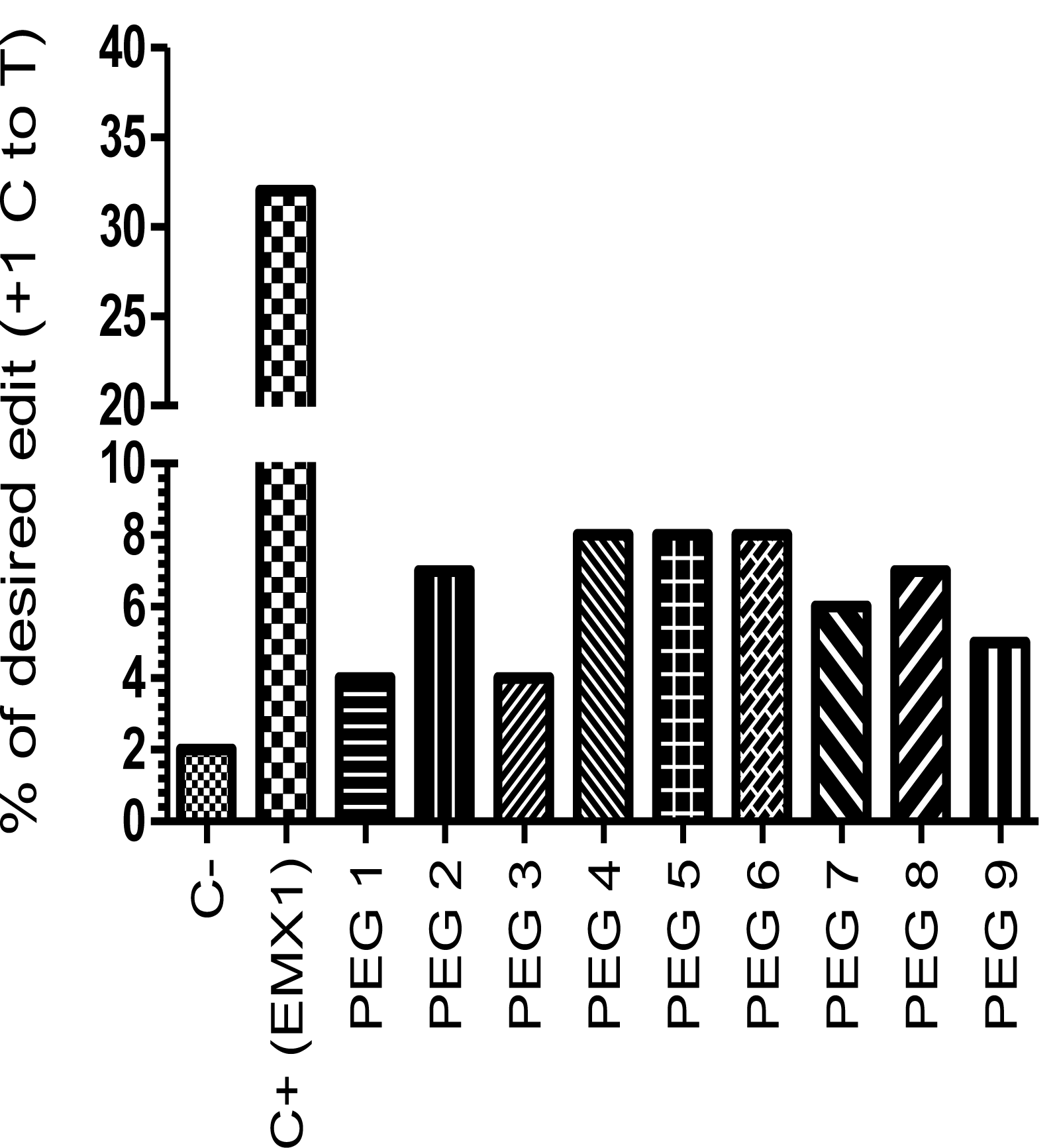
Prime editing of *DMD* exon 35. **A)** Partial sequence of exon 35 of the *DMD* gene. The SpCas9 PAM is in a red box and of the pegRNA protospacer sequence is underlined in red. **B)** The sequences of the Primer Binding Site (PBS, in bold) and of the Reverse Transcriptase Template (RTT, underlined) of 9 pegRNAs targeting *DMD* exon 35. The T in red in the RTT permits to introduce an adenosine (A) in the strand containing the PAM (in the present case the antisense strand). This results in the presence of a thymine (T) in the sense strand thus modifying the arginine codon into a stop codon. **C)** Results of Sanger sequencing and EditR analysis of PRIME editing of the *DMD* exon 35. There is a 2% background level of thymidine (T) in the position of the targeted nucleotide. However, several pegRNAs increased the percentage of Tup to 8%. A control pegRNA targeting the EMX1 gene produced a 32% modification in that gene.

We also designed 9 pegRNAs to induce a C to T mutation in a CGA arginine codon in *DMD* exon 35 to also induce a TGA stop codon (Figures 4A). Nine different pegRNAs were also tested (Figure 4B). Plasmids pU6-pegRNA-GG-acceptor coding for each of these pegRNAs under a U6 promoter were co-transfected in HEK293T cells with the pCMV-PE2 plasmid. Negative control cells were transfected with a plasmid containing a reporter eGFP gene. This permitted to confirm a good level of transfection. The DNA was extracted 3 days later and the mutations were analyzed by Sanger sequencing and the EditR web site. There was only 2% of T in the cytidine targeted site in the negative control cells. The percentage of C to T mutations in the targeted arginine codon ranged from 4 to 8% for the different pegRNAs. Given the presence of a 2% background, this mean up to 6% real targeted mutations.

Since the percentage of targeted mutations was low, we tested the hypothesis that it could be increased by multiple successive PRIME editing treatments (Figure 5A). We selected the 3 best pegRNAs identified in Figure 3C and 4C. For the first treatment (T1), HEK293T cells were transfected with pCMV-PE2 plasmid and plasmids coding for pegRNAs targeting the EMX1 gene or targeting exon 9 or exon 35 of the *DMD* gene. DNA was extracted from samples of the cells 3 and 6 days after the T1 treatment. Three days after this first treatment, the pegRNA targeting the EMX1 gene produced a 33% modification of the targeted nucleotide (Figure 5B). The pegRNAs targeting the *DMD* exon 9 produced only 9 to 14% modifications of the targeted cytidine into a thymine after this first treatment (Figure 5C). However, the Sanger sequencing noise was 8% in this experiment. For the pegRNAs targeting exon 35 of the *DMD* gene, the first PRIME editing treatment produced 5% to 8% modifications of the targeted C into a T (Figure 5D). However, the background noise was only 2% for this experiment. The percentages of mutations of the EMX1 and *DMD* gene remained relatively the same in samples obtained 6 days after the first treatment (Figures 5B, C and D). For the second treatment (T2), some cells from which the DNA was not extracted were transfected again the next day with the pCMV-PE2plasmid and a plasmid coding for the same pegRNA. DNA was extracted again 3 and 6 days after this second treatment (T2). The percentage of EMX1 mutations clearly increased to 49% (Figure 5B). The percentage of mutation of the targeted cytidine also increased with some pegRNAs (specially for pegRNA5 targeting *DMD* exon 35), which reached 14% 3 days after the second treatment (Figure 5D). However, the percentage of mutations induced by pegRNA2 targeting *DMD* exon 9 did not increase (Figure 5C). The mutagenic treatment was repeated a third time (T3) on the remaining cells and DNA was extracted again 3 and 6 days later. The percentage of mutations of the EMX1 did not increase further and remained at 49% (Figure 5B). However, the percentage of mutations of the *DMD* exon 35 increased again for pegRNA4 and pegRNA6 targeting *DMD* exon 35 reaching up to 16% (Figure 5D). The percentage of mutations induced by pegRNA2 targeting *DMD* exon 9 did not increase with repeated treatment.

**Figure 5).**
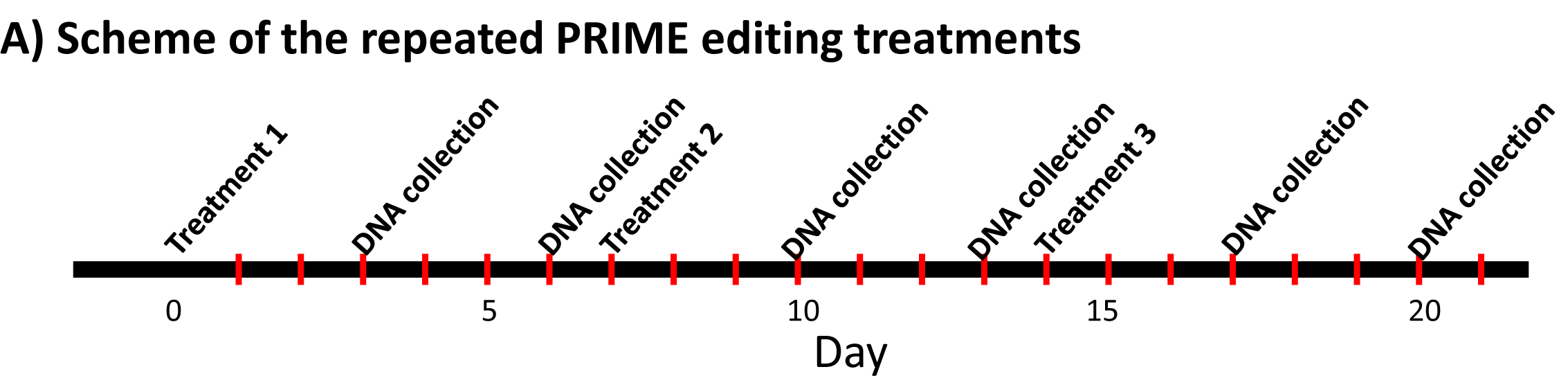
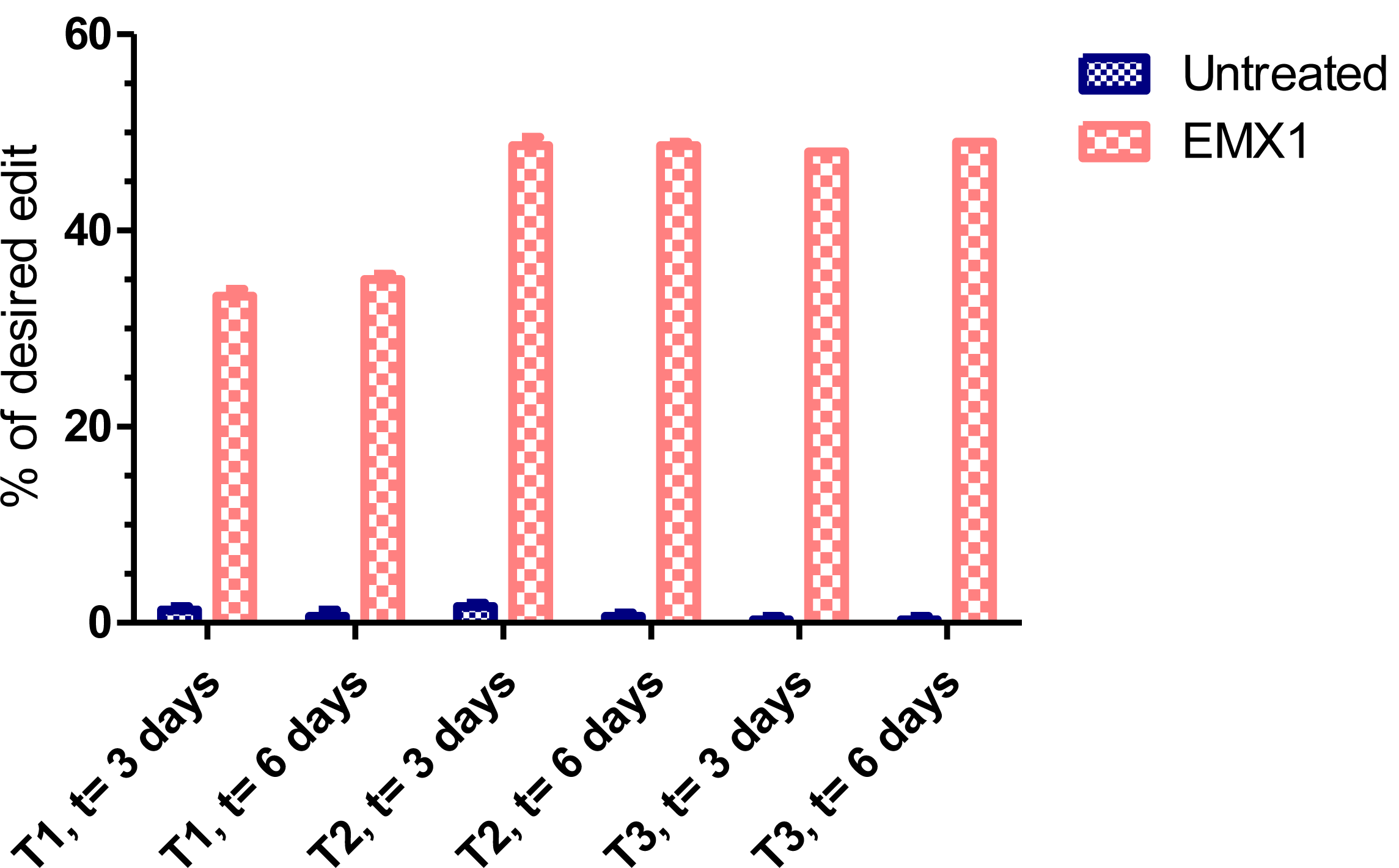
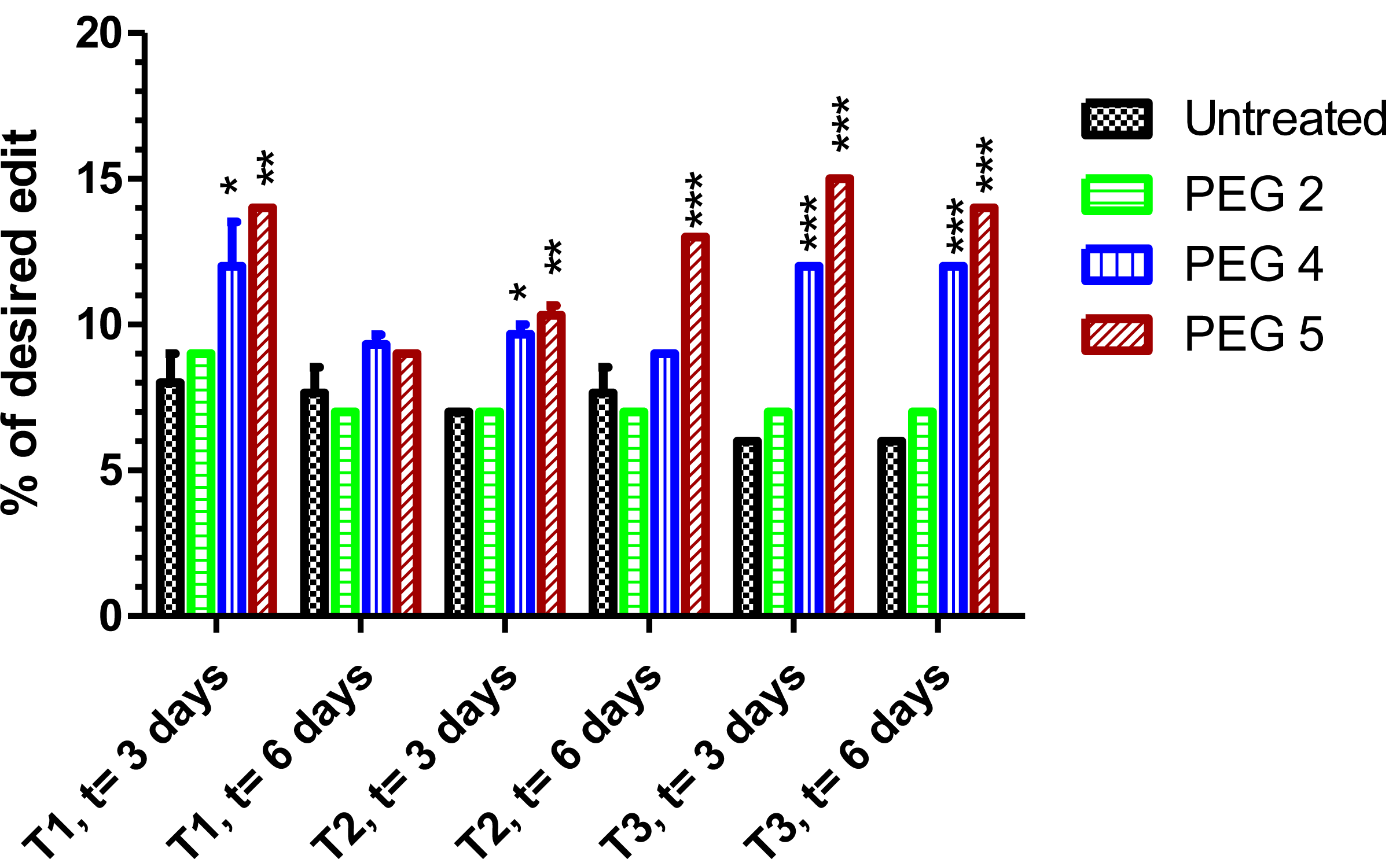
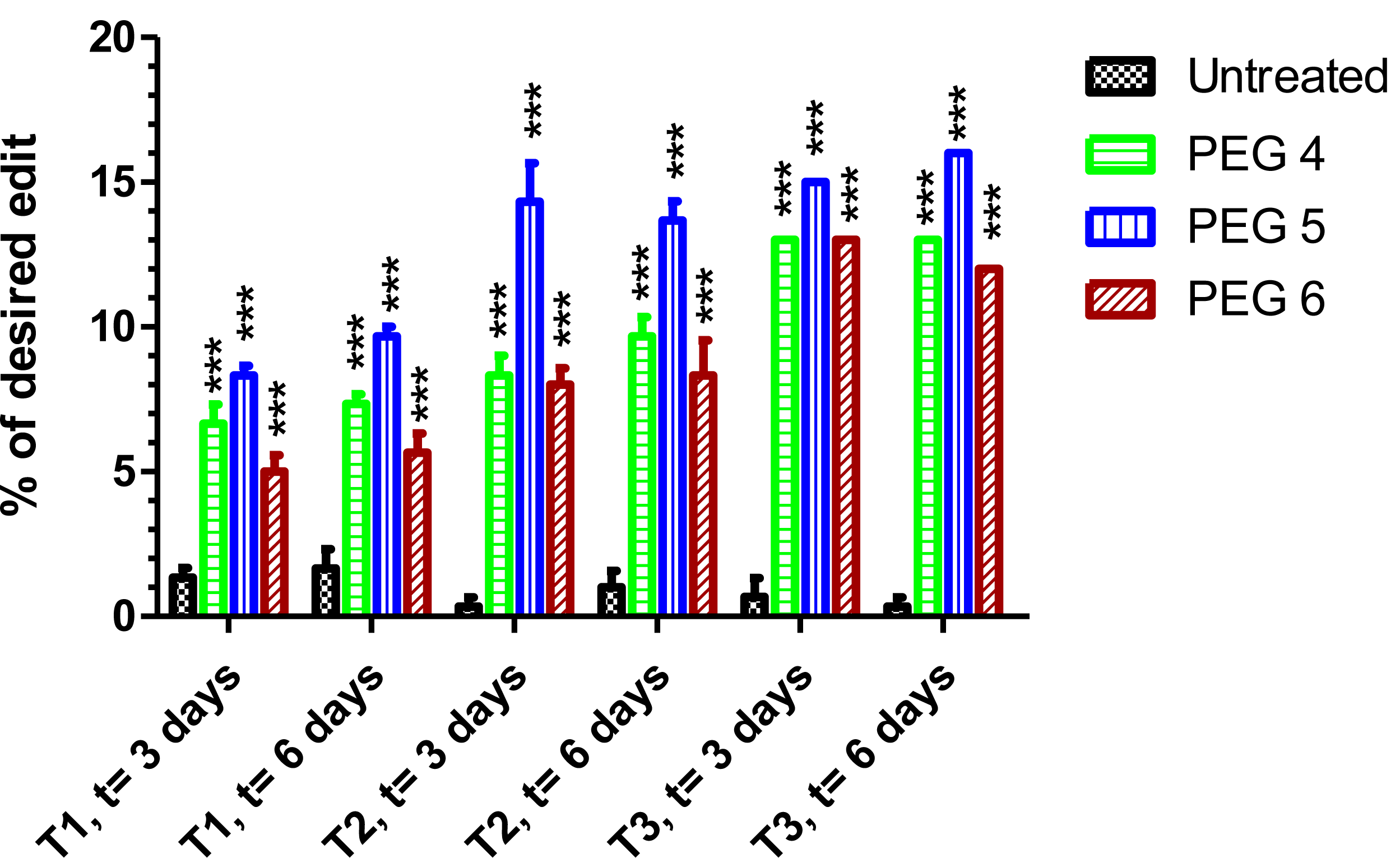
Repeated PRIME editing. **A)** Scheme summarizing the timing of the three successive PRIME editing treatments. The experiment was done in triplicates. HEK293T cells were transfected 3 times (at days 0, 7 and 14) with the plasmids pCMV-PE2 andpU6-pegRNA-GG-acceptor and with a second plasmid coding for a pegRNA under a U6 promoter. The pegRNAs were targeting either the EMX1 gene (**B**), exon 9 of the *DMD* gene (**C**) or exon 35 of the *DMD* gene (**D**). DNA was extracted from cell samples at 3 and 6 days after each treatment (i.e., at days 3, 6, 10, 13, 17 and 20). The percentages of desired nucleotide mutations were determined by Sanger sequencing and analysis with EditR. The figures illustrate the average and standard deviations. These percentages increased with the second treatment for the EMX1 gene and *DMD* exon 35. The third treatment with pegRNA5 and pegRNA6 increased the mutation of *DMD* exon 35. The results were analyzed by an analysis of variance. The *. ** and ***indicate respectively a p of less than 0.001, 0.0001. 0.00001 with the untreated controls.

## DISCUSSION

Our results confirm the results of Anzalone et al ^2^ that the PRIME editing technology may be used to induce specific nucleotide mutations. This is very encouraging for the possibility of using this technique for the correction of point mutations responsible for hereditary diseases. However, our experiments were done in a cell line and thus the cells were proliferating. Thus, in future experiments, we will have to confirm that specific nucleotide mutation may be induced *in vivo* directly in the cells that do not proliferate. This will have to be tested in neurons for Alzheimer’s disease mouse models and muscle fibers for Duchenne Muscular Dystrophy (DMD) mouse models.

The fact that the percentage of nucleotide mutations increased with repeated PRIME editing treatment is very encouraging. It is also important to note that up to 15% nucleotide mutations were obtained in the *DMD* gene. Muscle fibers contain thousands of nuclei (i.e., up to 60 nuclei par mm).The nuclear domain^7^ (i.e., the length of muscle fiber membrane over which the dystrophin is present when there is only one competent nucleus is about 439 μm^8^. Since there are about 30 nuclei in the fragment of a muscle fiber, this means that the modification of the *DMD* gene in 1 out of 30 nuclei, i.e., only 3% of the DMD genes, would be sufficient for the expression of dystrophin over most of the muscle fiber membrane and thus sufficient to produce a therapeutic effect. Thus, the percentage of nucleotide modifications induced by the PRIME editing technique is above the minimum required to expect a beneficial outcome, if such a percentage may be obtained in muscle fibers.

It is clear that for the PRIME editing technique, several different pegRNAs have to be tested, since some pegRNAs did not work at all. Additional research will have to be done to understand the reason of poor and good results to improve the design of the pegRNAs. Moreover, some genes like the EMX1 seems to be more easily edited. The repeated administration of some pegRNAs did not increase the percentage of gene correction while there was good improvement for other pegRNAs, we have to understand why. The percentage of specific nucleotide mutations in the EMX1 gene also increased with a second administration reaching 49%, however the percentage of corrections did not increase further with a third treatment. Again, this phenomenon has to be understood.

The percentage of modification of the *APP* gene to introduce the A673T mutation is however only 5% with the best pegRNA (pegRNA1). This is too low to expect an eventual therapeutic effect. This the PRIME editing technique has to be improved for Alzheimer treatment and probably for many other hereditary diseases, which will also require a high percentage of modification of the target gene to obtain a therapeutic effect.

## MATERIAL AND METHODS

### Plasmid

Prime Editor 2 (PE2) plasmid coding for the SpCas9 gene fused with the MLV reverse transcriptase was obtained from Addgene Inc. (pCMV-PE2 # 132775). The plasmid used to express a pegRNA was also obtained from Addgene Inc. (pU6-pegRNA-GG-acceptor # 132777). Cloning of pegRNA into the BSAI-digested pU6-pegRNA-GG-acceptor was as described by Anzalone and al. ^2^. Oligonucleotides used to construct all pegRNAs were obtained from IDT inc.

### Cell culture

HEK293T were grow in DMEM-HG medium (Wisent Inc.) supplemented with 10% FBS (Wisent Inc.) and 1% penicillin-streptomycin (Wisent Inc.) at 37 °C and with 5% CO_2_, in a humidified incubator. The day before transfection, cells were detached from the flask with a Trypsin-EDTA solution (Sigma Inc.) and counted. Cells were plated in a 24 wells plate at a density of 60 000 cells per well with 1 ml of culture medium. The day of the transfection, medium was replaced with 500 μl of fresh medium. Cells were transfected with 1 μg of total DNA (500 ng of each plasmid when co-transfection was required) with lipofectamine 2000 according to the manufacturer instruction. The day after, medium was changed for 1 ml of fresh medium and cells were maintained in incubator for 72 hrs before genomic DNA extraction. For multiple treatment experiments, cells were plated on day 0 and transfected as described. On day 3, 50% of cells were recovered for genomic DNA preparation and the 50% left were plated in one well of a 6 wells plate and let them grow until the day 6. Cells were detached from the well with trypsin-EDTA solution, counted and DNA was extracted from a sample of the cells. The remaining cells were plated at 60 000 cells per well in a 24 wells plate. The following day, cells were transfected as described. We used the same procedure for the third treatment.

### Genomic DNA preparation and PCR amplification

Cells were detached from wells directly with up and down pipetting in the culture medium and transferred in Eppendorf tubes. Cells were spin 5 minutes at 9000 RPM on microcentrifuge at room temperature. Cell pellets were washed once with 1 ml of PBS 1X and spin again for 5 minutes at 9000 RPM. Genomics DNA were prepared using the DirectPCR Lysis Reagent (Viagen Biotech Inc.). Briefly, 50 μl of DirectPCR Lysis Reagent containing 0.5 μl of a proteinase K solution 20 (mg/ml) were added to each cell pellet and incubated overnight at 56°C followed by another incubation at 85°C for 45 minutes. One μl of each genomic DNA preparation was used for the PCR reaction. For each primer sets (Table 1) 50 μ l of PCR reactions was as follows: 98°C: 30 seconds, (98°C: 10 seconds, 60°C: 20 seconds, 72°C: 45 seconds) for 35 cycles. A final extension at 72°C for 5 minutes was also performed. We used Phusion ™High-Fidelity DNA polymerase from Thermo Scientific Inc. for all PCR reactions. 5 μl were electrophorized in 1X TBE buffer on 1 % agarose gel to control the PCR reaction qualities and to make sure that only one specific band was present. The 45 μl PCR reaction left were Sanger sequenced with an internal primer (Table 1). Sequencing results were analyzed with the EditR web site (https://moriaritylab.shinyapps.io/editr_v10/).

### Deep sequencing analysis

Deep sequencing samples were prepared by a PCR reaction (as described) with special primers containing a bar code sequence (BCS) sequence to permit the subsequent sequencing (Table 1). PCR samples were sent to Centre d’Innovation Génome Québec at McGill University sequencing facilities to sequence amplicons with an Illumina sequencer. Roughly 5000 to 10000 reads were obtained per sample.

### Bioinformatics analysis of the deep sequencing results

The proportion of wild-type versus edited genomes was estimated by counting the abundance of sequenced reads that contained exon 16 with the C wild-type codons and the T edited codons.

## Statistical analysis

Data were analysed using the Graph Pad PRISM 5.0 software package (Graph Pad Software Inc., La Jolla, California, USA). The mean between different PEGs and negative control were compared using the two ways ANOVA repeated measures. A p value less than 0.05 were considered statistically significant for a 95% confidence interval.

## Acknowledgments

This work was supported by grants from Weston Brain Institute, Jesse’s Journey the Foundation for Cell and Gene Therapy and the Canadian Institute of Health Research (CIHR). AG has a studentship from the CHUQ foundation and CM has a studentship from Centre Thématique de Recherche en Neurosciences (CTRN).

## Author Contributions

JR, CHM, GT and FGB did the experiments, AG analyzed the deep sequencing results and JPT, JR and CHM planned the experiments and wrote the manuscript.

## References

1 Komor, A. C., Kim, Y. B., Packer, M. S., Zuris, J. A. & Liu, D. R. Programmable editing of a target base in genomic DNA without double-stranded DNA cleavage. Nature 533, 420–424, doi:10.1038/nature17946 (2016).

2 Anzalone, A. V. et al. Search-and-replace genome editing without double-strand breaks or donor DNA. Nature, doi:10.1038/s41586-019-1711-4 (2019).

3 Goate, A. et al. Segregation of a missense mutation in the amyloid precursor protein gene with familial Alzheimer’s disease. Nature 349, 704–706, doi:10.1038/349704a0 (1991).

4 Jonsson, T. et al. A mutation in APP protects against Alzheimer’s disease and age-related cognitive decline. Nature 488, 96–99, doi:10.1038/nature11283 (2012).

5 Roberts, R. G., Gardner, R. J. & Bobrow, M. Searching for the 1 in 2,400,000: a review of dystrophin gene point mutations. Hum Mutat 4, 1–11 (1994).

6 Kleinstiver, B. P. et al. Engineered CRISPR-Cas9 nucleases with altered PAM specificities. Nature 523, 481–485, doi:10.1038/nature14592 (2015).

7 Blau, H. M., Pavlath, G. K., Rich, K. & Webster, S. G. Localization of muscle gene products in nuclear domains: does this constitute a problem for myoblast therapy? Adv Exp Med Biol 280, 167–172 (1990).

8 Kinoshita, I., Vilquin, J. T., Asselin, I., Chamberlain, J. & Tremblay, J. P. Transplantation of myoblasts from a transgenic mouse overexpressing dystrophin produced only a relatively small increase of dystrophin-positive membrane. Muscle Nerve 21, 91–103 (1998).

